# The interplay of immune components and ECM in oral cancer

**DOI:** 10.1101/259622

**Authors:** Yigang Kwak, Burair Alsaihati, Chi Zhang, Ying Xu, Sha Cao

## Abstract

We report that in Oral Squamous Cell Carcinoma (OSCC), extracellular matrix (ECM) plays a vitally important role in defining the characteristics of cancer vs. normal, as it is a compartment with significant enrichment of known OSCC biomarkers, and the number of genes constituting ECM are more prominently upregulated in OSCC than almost all the rest of the cancer types. This is probably due to the constant exposure of oral cavity to external stimuli, resulting in the ECM remodeling, which further is a key player in tumor invasion. While we showed ECM molecules alone could well distinguish oral cancer from normal tissue samples, a significant portion of these predictive ECM molecules, share the same transcriptional regulator, NFKB1, a master regulator of immune response. We further studied the level of involvement of the immune system in OSCC, and found that the immune composition in OSCC is distinctly different from the other cancer types. OSCC has a higher level of infiltration of adaptive immune cells, including B cell, T cell and neutrophil, compared with other cancer types, while a lower level of infiltration of innate immune, including macrophage and monocyte. Previous studies have revealed the roles of ECM and immune system in OSCC development, and our study showed that ECM plays a very prominent role in OSCC, subject to the complex microenvironment in oral cavity, particularly the immune system profile, and our association analysis revealed it is likely the interactions between ECM and immune cells that define the highly invasive property of OSCC.

## Introduction

Squamous Cell Carcinoma (SCC) is the most common malignant neoplasm of the oral cavity, which represents around 90% of all oral cancers [1], which puts OSCC the sixth most widespread malignancy with an average 5-year survival rate of approximately 60% [2, 3]. Several risk factors have been associated with increased risk of developing oral cancer, most notably tobacco use and heavy alcohol consumption. Tobacco contains pre-carcinogenic elements that promote cancer, and these elements undergo coordinated alterations by oxidative enzymes and result in electron-poor products and agents that covalently bind to the DNA, producing adduct mutated regions [4]. In addition, consumption of alcohol increases risk of oral cancer development because alcohol causes epithelial atrophy and inhibit DNA synthesis and repair from the dissolvement of lipid in epithelium and increased permeability of oral mucosa. Likewise, alcohol is known cause systemic effects, especially in immunosuppression. The chronic use of alcohol is known to impair both the innate and acquired immunity, which escalates susceptibility to infection and neoplasms. [5]. Furthermore, substantial evidence suggests that a large array of microbial species can encourage the initiation, promotion, or progression of human malignances. Especially because the mouth comprises of more than 750 oral bacterial taxa, oral epithelium is prone to constant microbial challenges, which can lead to malignances.[6, 7].

The tumor microenvironment of OSCC is genetically heterogenous, but a several common grounds to its TME is depicted in the following. For instance, many discoveries in molecular oncology has shown that the progression of dysplastic lesions to malignant forms include cellular elements that are not only tumor cells, but also other nonmalignant cell types, such as innate and adaptive immune cells, fibroblasts, epithelial cells and endothelial cells. [8]. These cellular elements of tumor microenvironment often coevolve with the tumor in a disorganized response. Namely, the tumor invasiveness is increased by release factors such as matrix metalloproteins (MMPs), which are from cancer-associated fibroblasts [9, 10]. In addition, when the chronic inflammation of TME is unresolved, numerous changes in adaptive immune response occurs, such as apoptosis of cytotoxic T cells and activation of suppressor T cells [10]. Furthermore, the famous “seed and soil” hypothesis applies where tumors are capable of reprogramming their microenvironment to a metabolically fertile environment that is susceptible for their high energy and anabolic requirements [11]. Lastly, many critical targets, such as NF-kb, hypoxia-inducible factor-1alpha, and VEGF have been continued to be explored as therapeutic targets in the tumor microenvironment [12–14].

Tumor invasion in OSCC is caused by the three-step process that involves changes in tumor cell adhesion, degradation of collection of molecules, such as basement membranes, extracellular matrix (ECM), and matrix metalloproteases, and migration of tumor cells in proteolytically modified ECM [15, 16]. In most epithelial layers, including the oral cavity, the extracellular matrix is composed of laminin, collagen, fibronectin, and glycosaminoglycans distinctively from keratinocytes and stromal cells, or from concerted interactions between these two cell types. The function of ECM is known to be significant for tissue development, adult tissue maintenance, wound healing, and oncogenesis[17]. The oncogenic and tumor development is encouraged when the ECM composition changes by altered expression and secretion. In some cases, it has been reported that further changes in ECM receptors are imperative for converting premalignant squamous epithelium to malignant lesions. Also, increased collagen type I expression is reported in malignant transformation of keratinocytes and associated with well-differentiated OSCC [18], and changes in collagen expression promote adhesion, migration, and differentiation. In addition, collagen VII and XVII seem to play a role in OSCC development and progression [15].

Despite the availability of established OSCC biomarkers and the prevalent evidence of ECM role in most cancers, the extent of ECM significance in the specific context of OSCC has not been thoroughly investigated, and the knowledge about its interaction with the OSCC microenvironment and its biomarkers is limited. This knowledge is crucial to understand the microenvironment changes in OSCC and set the compass for future research aimed at searching for OSCC-specific targeting therapies. In this work, we show that ECM components and features play a central role in OSCC development, far beyond their involvement in many other cancers. We shed some light on ECM features that have consistent overexpression levels in OSCC tissues, immune cells with increased and decreased population in OSCC and the ECM components that interact with each immune system component.

## Materials and Methods

### Transcriptomic data collection and normalization

Gene expression data were obtained from GEO and TCGA databases. We initially collected 27 GEO datasets that contain oral cancer samples, and then filtered out those datasets where less than 5 normal or cancer samples are available, and only 10 datasets were left in the study. These microarray-based gene expression data were all normalized using MAS5. The TCGA Head-Neck Squamous cell Carcinoma (HNSC) expression dataset contains large portion of oral samples. All the TCGA expression data are RNA-Seq based, with FPKM normalized gene expression quantification with further upper-quantile normalization. The following table describes each dataset and the associated sample size.

### Immune cell deconvolution

The prediction of the immune composition of tumor tissue samples was performed using CIBERSORT [19]. For GEO datasets, probe ids were converted to Hugo gene symbols first. Probes with no corresponding genes were removed and genes with multiple probe ids were mapped to the probe with the highest mean expression level across all samples within its dataset. For TCGA-HNSC dataset, Ensembl transcript ids were mapped to Hugo gene symbols. Transcripts without known gene names were removed. For each dataset, whether from GEO or TCGA, genes with missing values in any sample were further removed before Cibersort analysis. Finally, for each dataset, all the remaining gene expression values were passed to CIBERSORT R function (release v1.03) using LM22 as a signature file, number of permutations equal to 500 and quantile normalization set to true [19].

The original paper suggested the method works best with MAS5 normalized expression data, so our deconvolution analysis was only conducted with those datasets on Affymetrix platforms, with.CEL formatted raw data available.

### Differential expression and enrichment analysis

Differential expression analysis was performed using t test, with significance threshold 0.001. Enrichment analysis was performed using hypergeometric test, with online database DAVID [20].

### ECM gene lists

The ECM genes are collected from pathways involved in ECM composition, organization, assembly and disassembly, altogether 123 genes. Specifically, the ECM genes we used contain genes from the following gene sets from Msigdb [21]: REACTOME DEGRADATION OF THE EXTRACELLULAR MATRIX; REACTOME EXTRACELLULAR MATRIX ORGANIZATION; GO REGULATION OF EXTRACELLULAR MATRIX ORGANIZATION; GO REGULATION OF EXTRACELLULAR MATRIX ASSEMBLY; GO POSITIVE REGULATION OF EXTRACELLULAR MATRIX ORGANIZATION; GO REGULATION OF EXTRACELLULAR MATRIX DISASSEMBLY.

### Transcription factor and pathway databases

Experimentally validated human transcription factors (TFs) and their target genes were collected from multiple databases: TRED [22], Neph2012 [23], ENCODE [24], Marbach2016 [25]and TRRUST [26]. Overall, 919 transcription factors were retrieved. A TF-target relationship is considered as reliable only when the relationship appears in at least two of these databases. Overall, 72,407 pairs of such relationships were collected and used in our analyses.

### Penalized logistic regression

For each response variable, we collected the corresponding predictors as the linear model predictors. We have built a logistic regression model with Li penalty to select the most parsimonious subset of predictors that minimizes the following objective function:

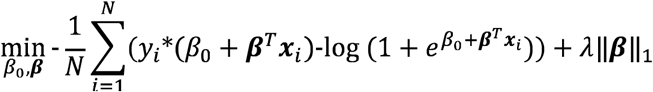

where *y*_*i*_, ***X***_***i***_ are the *i*th observed response and predictor, respectively; *β_0_,* ***β*** are coefficients; and A is the overall penalty parameter, which is selected based on cross-validation performance. In the case when the number of predictors are fixed, we select A so that the selected predictors (fixed number) can explain the highest percentage of deviance.

## Results

### ECM involvement in harboring biomarkers in oral cancer

We comprehensively collected all the cancer biomarker genes for OSCC from large amounts of literature in PubMed database, and the full list contains 75 protein coding genes, available in Supplementary Table S1. These biomarkers were collected because there are research papers in PubMed database that reported their gene products are significantly up-regulated over oral cancer development stage. We looked up the sub-localizations of these molecules, among nucleus, extracellular space, Golgi, cytosol, ER, and noticed that 37 out of 75 biomarkers are located in the extracellular space, 12 out of 75 in the extracellular matrix. Gene set enrichment analysis using DAVID [20] showed significant enrichment of these biomarkers in extracellular space (3.50E-20) and ECM (8.40E-07). (see Supplementary Table S2). These biomarkers clearly bear strong biological significance of oral cancer development, and in order to derive some mechanistic interpretation of these biomarkers, and further of OSCC, we aim to first select those biomarkers that indeed show robust and stable up-regulation in oral cancer vs. normal.

We collected 10 sets of gene expression data on different high-throughput microarray platforms, and 1 set of RNA-Seq based high-throughput gene expression data from The Cancer Genome Atlas (TCGA) database (see Table 1). We analyzed differential expressions for each of the molecules using hypothesis testing (see Methods and Materials) for each dataset, and while these genes are expected to show significant up-regulation in cancer vs. normal samples, a large portion didn’t appear this way (Figure 1A). This could be resulted from the fast degradation rates of some mRNAs that caused its loss of abundance. We observed that molecules consistently significantly up-regulated in cancer patients’ compared with normal across in at least eight dataset are: CXCL1, EGFR, GREM1, IFI44, IFI6, INHBA, ISG15, ITGA3, LAMC2, LDHA, LGALS3BP, MICB, MMP1, MMP10, MMP3, MMP9, MYH9, PLAU, POSTN, PTHLH, TNC. 12 out of 21 biomarkers are located in the extracellular space, 6 out of 21 in the extracellular matrix. Gene set enrichment analysis using DAVID showed significant enrichment of these significant biomarkers in extracellular space (2.20E-06) and ECM (2.50E-04) (see Supplementary Table S3). Notably that four MMP (matrix metalloproteinase) genes are found consistently up-regulated in OSCC cancer. MMPs degrade basement membranes and ECM, reported to be instrumental in OSCC development [27]. Here we demonstrated that a significant portion of the biomarkers reside in ECM, and among those robust biomarkers, ECM again accounts for a significant portion.

**Table 1:** The columns are: (1)&(2) data source and accession number; sample sizes for (3) normal and (4) cancerous expression data. Note that GSE2109 has multiple cancer types, and was used to compare the immune cell relative proportions across multiple cancer types.

| Source | Dataset | Normal samples | Tumor samples |
| --- | --- | --- | --- |
| GEO | GSE13601 | 26 | 31 |
|  | GSE34105 | 16 | 62 |
|  | GSE34106 | 15 | 28 |
|  | GSE46802 | 9 | 9 |
|  | GSE23558 | 5 | 27 |
|  | GSE25099 | 22 | 57 |
|  | GSE25104 | 22 | 57 |
|  | GSE30784 | 45 | 167 |
|  | GSE37991 | 40 | 40 |
|  | GSE38532 | 40 | 40 |
|  | GSE2109 | 0 | multiple |
| TCGA | OSCC/HNSC | 32/44 | 379/520 |
|  | BLCA | 19 | 408 |
|  | BRCA | 113 | 1095 |
|  | COAD | 41 | 285 |
|  | KIRP | 32 | 290 |
|  | LIHC | 50 | 371 |
|  | LUAD | 59 | 515 |
|  | LUSC | 51 | 501 |
|  | PRAD | 52 | 497 |
|  | THCA | 59 | 505 |

**Figure 1:**
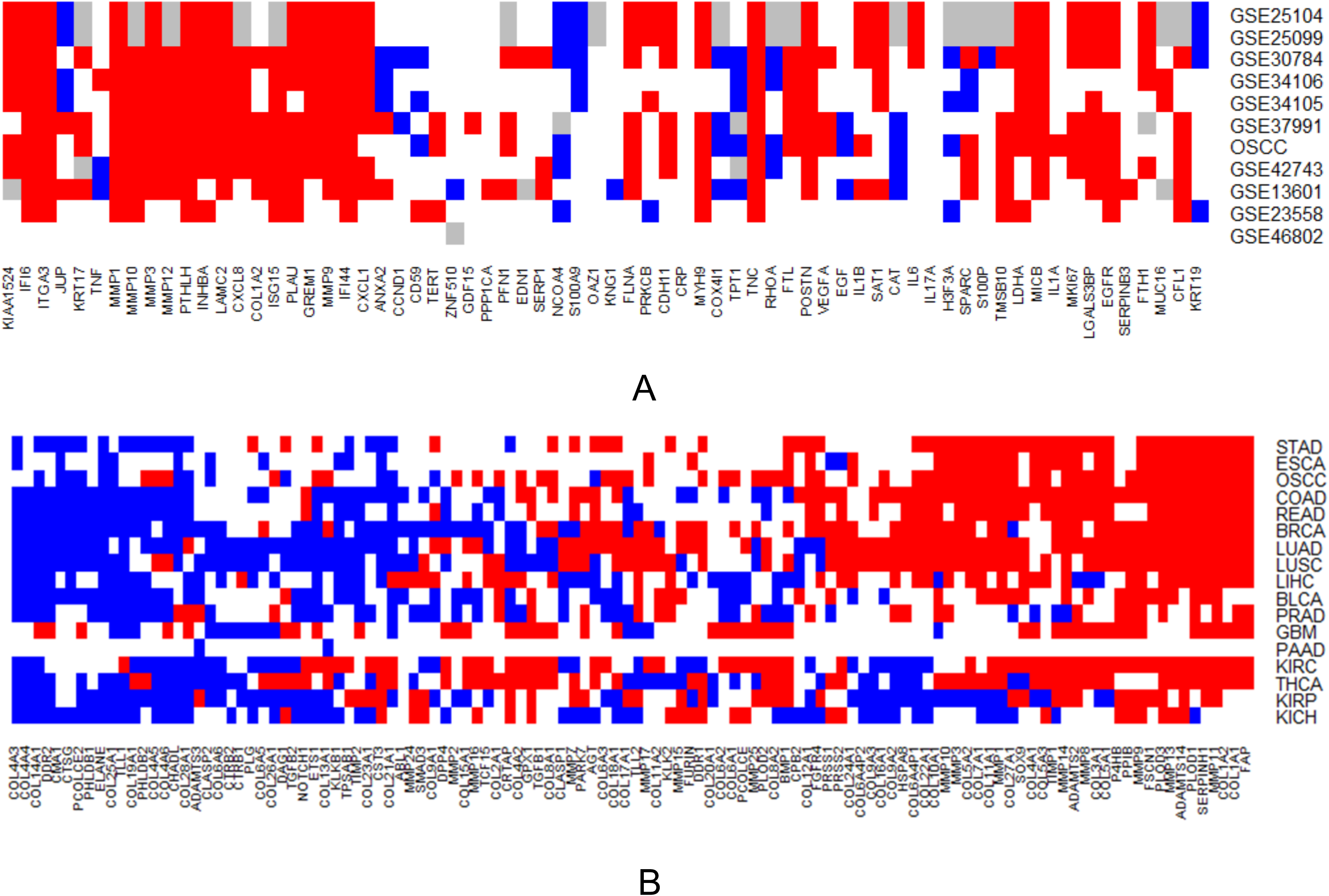
A) Differential expressions of the reported biomarkers of OSCC; B) Differential expressions of ECM molecules across multiple cancer types. *x*-axis are genes, *y*-axis are datasets or cancer types. Red cubes indicates significant up-regulation of genes in cancer with *p*-value <0.01 and blue indicates significant down-regulation with *p*-value <0.01, gray cubes indicate there is a lack of information, and white cubes mean that the genes are not significantly different between tumor and normal samples.

### ECM markers are good predictors of OSCC cancer

In light of the significance of ECM in fostering OSCC biomarkers, we collected large number of genes involved in ECM constituents, and its organization, which includes ~100 genes that covers a wide range of ECM functional entities (See supplementary Table S4). We noticed that among all the cancer types, LUAD, KIRC and OSCC, seems to have the most number of ECM genes, which are 58, 56, 56, that are significantly up-regulated than their normal counterpart (Figure 1B). This reflects the high level specificity of ECM in OSCC cancer development, in addition to LUAD and KIRC. We further extracted the expression values of the ECM genes, in both tumor and normal samples of each oral cancer dataset collected. We then built a penalized logistic regression model, with the response variable being a binary variable indicating whether the tissue sample is tumor (=1) or normal (=0), and the ECM expression values are used as linear predictors. The penalty parameter λ was selected by 10-fold cross-validation that achieves the lowest average classification error on the training data sets. We also did the same analysis using the whole set of genes, trying to see whether the ECM gene set alone could predict OSCC tumor and normal class just as well as using all the genes. The deviance explained using merely ECM molecules and the whole gene set are listed in Table 2, which appear to be very close. Notably, MMP1 has been selected as a potential predictor of OSCC cancer vs. normal by penalized logistic regression model for all the datasets being considered. We collected those genes that are both significantly up-regulated in five datasets, and have been selected by penalized logistic regression model (using whole gene set) in at least three datasets, and they include: *COL4A1, COL4A2, COL4A5, COL4A6, COL7A1, MMP1, MMP3, MMP9, MMP11, MMP13, PLOD1, SERPINH1, FSCN1, FAP.* Interestingly, 5 out of the 14 genes, *MMP9, MMP3, COL7A1, MMP1, MMP13,* share the same transcription factor, *NFKB1,* which is a master regulator of immune responses [28]. This drives us to further study the interplay between ECM and immune system.

**Table 2:**
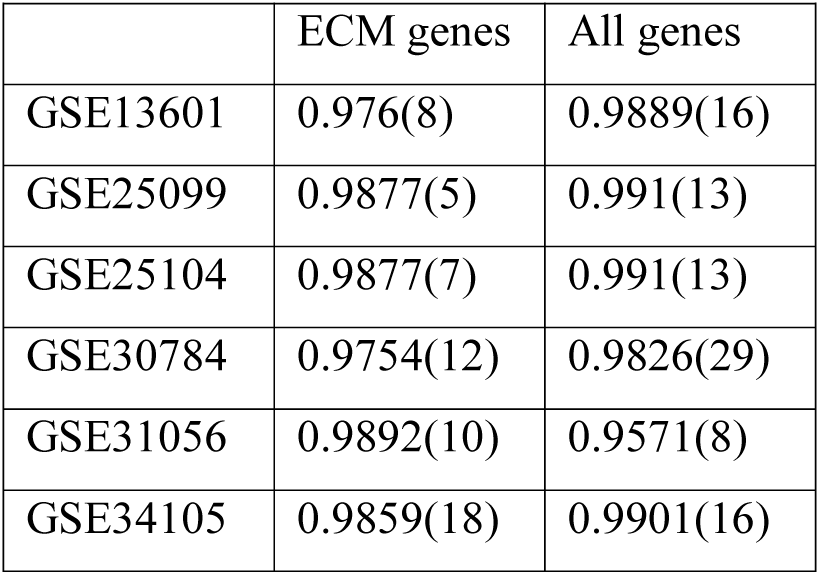
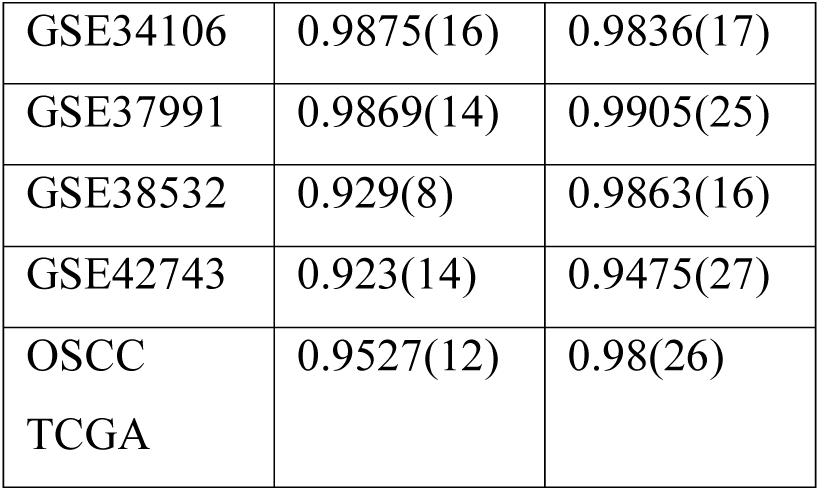
the deviance explained by using ECM genes only (column 2) and whole gene set (column 3). The number in the parenthesis is the number of genes selected as predictors.

**Table 3:** the ECM genes that are shown to be strongly associated with immune cell types in at least 5 datasets..

|  |  |
| --- | --- |
| B | SMAD3,MMP13,COL1A2,PLOD3,COL6A3 |
| Plasma | SERPINH1,PLOD1,COL5A2,MMP1,FSCN1 |
| T | DDR2,PLOD2,CMA1,COL18A1,TIMP1,SERPINH1,MMP14,COL6A3,MMP9,COL5A2<br>,COL16A1,MMP11,CMA1,P4HB,FSCN1,MMP7 |
| NK | COL4A1 |
| Monocytes | MMP10 |
| Macrophages | PLOD1,COL6A3,MMP11,SOX9,COL17A1,COL18A1,COL5A2,PLOD3,COL6A3,MMP<br>24,MMP1,BMP1,COL5A2 |
| Dendritic | SMAD3,MMP9,TIMP1,MMP11,MMP7 |
| Mast | SERPINH1,TIMP2,COL5A2 |
| Eosinophils | COL5A2,COL8A1,MMP1 |
| Neutrophils | MMP7 |

### Significant different immune cell population in OSCC

Knowing the significant associations between ECM components and the immune system and to thoroughly examine the interactions between the two, we used the state of the art deconvolution method to estimate the relative immune cell populations in OSCC, as well as a multitude of other cancer types. The essence of the deconvolution method is that, some genes, commonly called marker genes, are known to be essentially expressed only in one or a few specific cell types, and estimating cell type proportions is based on the expression signature of these marker genes in the tissue mixture. We noticed that the following cell types are significantly less in OSCC than other cell types: macrophage and monocyte (Figure 2). Compared with other cancer types, OSCC seems to have relatively more prominent amount of B cell, T cell and neutrophil, than the rest of the cancer types (Figure 2). In summary, OSCC has relative higher infiltration of adaptive immune cells, lower infiltration of innate immune cells. The oral cavity acts as the gateway for both the gastrointestinal and respiratory tracts, its immune environment must be equipped to prevent pathogen entry while maintaining immune homeostasis, which may be achieved by a very different way from the immune system of other tissue types [29].

**Figure 2:**
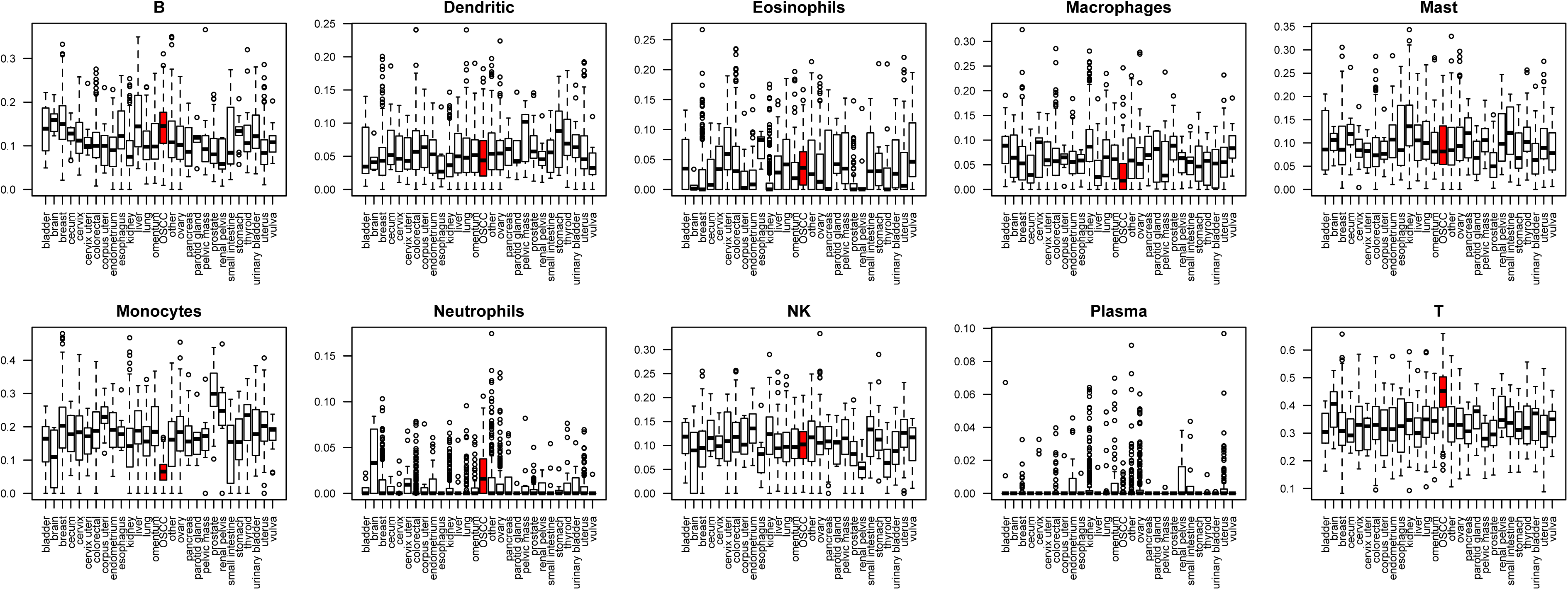
The relative proportions (*y*-axis) of different immune cell types in different cancer types (*x*-axis). Each panel shows the relative proportions of one immune cell type across multiple cancer types. The proportions for OSCC are marked by red bars.

We did co-expression analysis using the immune cell relative proportions with the ECM component genes. We speculate that immune cell infiltration in the tumor tissue could cause the remodeling of ECM by releasing relevant molecules that change the structure of ECM. Previous studies have shown that the inflammatory microenvironment surrounding the tumor is known to be involved in the induction, selection, and expansion of malignant cells [30]. Inflammation can have several effects on cancer, as acute inflammation has been said to counteract cancer, whereas, chronic inflammation has been seen to promote cancer development. In the event of chronic inflammation cascade, cytokines, chemokines, prostaglandins and reactive oxygen and nitrogen radicals accumulate in the microenvironment of tissues affected by chronic inflammation. If persistent, these inflammatory factors induce cell proliferation and promote prolonged cell survival through activation of oncogenes and inactivation of tumor-suppressor genes. This may result in genetic instability with an increased risk of cancer [31, 32]. Increased cell proliferation and cell survival may also bring about the production and secretion of inflammatory mediators. These biological mediators generate an inflammatory microenvironment that further increase cell survival and proliferation of the transformed cells, as well as promoting angiogenesis and evasion of protective immune responses [31, 33]. Once an inflammatory microenvironment has been established, reciprocal interactions between the evolving tumor cells and their stromal cells sustain cancer cell proliferation and promote progression of the tumor[34, 35].

## Discussion and conclusions

In light of the strong implications of microenvironment on oral cancer development, we presented a computational study on the roles of ECM, stromal and immune cells on OSCC development. It turns out that ECM is the compartment where the most significant portion of cancer specific events happen, which means that most of the OSCC biomarkers are found to be located in ECM. In a study for metastatic distribution acquired from Harvard, Palmer, Pondville, and Westfield medical centers [36], where the frequency of 30 metastatic sites from 41 primary sites were calculated based on 9487 incidences, oral cancer, specifically tongue cancer, has a very high tendency of metastasis. Metastasis is a process where neoplastic cells must be capable of degrading the ECM, largely through MMPs. Our results, as well as other studies, have shown that MMPs indeed play an important role in oral cancer [27]. Even though it wasn’t clear whether the invasion system through ECM is particularly favored by oral cancer, we showed that in oral cancer, compared with majority of other cancer types, indeed have more involvement of ECM in its development, as shown by the higher number of ECM genes significantly up-regulated in OSCC than the majority of other cancer types. The fact that a few significantly up-regulated ECM molecules share the same regulator as immune system, NFKB1, suggesting the link of these two systems. And the invasiveness of OSCC induced by ECM remodeling is probably accompanied by the local microenvironment, especially the immune system.

The current study is a systematic evaluation of the ECM and immune system in OSCC using multiple gene expression data, and the findings are believed to be consistent and robust for multiple datasets. Due to the limitations of interpretations by gene expression data, further experimental validation is required, and more thorough understanding of the interactions between ECM and immune cells in OSCC will provide an opportunity to refine the therapeutic interventions that are currently undertaken.

## Author contributions

S Cao conceived and designed the project; J Kwak, S Cao, and B Alsaihati all performed the analysis; C Zhang and Y Xu helped writing the article.

## Acknowledgement

The authors would like thank Ms. Xu Zhang for her help in collecting oral cancer expression data sets.

